# Universal nature of drug treatment responses in drug-tissue-wide model-animal experiments using tensor decomposition-based unsupervised feature extraction

**DOI:** 10.1101/2020.03.08.982405

**Authors:** Y-h. Taguchi, Turki Turki

## Abstract

Gene expression profiles of tissues treated with drugs have recently been used to infer clinical outcomes. Although this method is often successful from the application point of view, gene expression altered by drugs is rarely analyzed in detail, because of the extremely large number of genes involved. Here, we applied tensor decomposition (TD)-based unsupervised feature extraction (FE) to the gene expression profiles of 24 mouse tissues treated with 15 drugs. TD-based unsupervised FE enabled identification of the common effects of 15 drugs including an interesting universal feature: these drugs affect genes in a gene-group-wide manner and were dependent on three tissue types (neuronal, muscular, and gastroenterological). For each tissue group, TD-based unsupervised FE enabled identification of a few tens to a few hundreds of genes affected by the drug treatment. These genes are distinctly expressed between drug treatments and controls as well as between tissues in individual tissue groups and other tissues. We also validated the assignment of genes to individual tissue groups using multiple enrichment analyses. We conclude that TD-based unsupervised FE is a promising method for integrated analysis of gene expression profiles from multiple tissues treated with multiple drugs in a completely unsupervised manner.

## BACKGROUND

Drug design is a time-consuming and expensive process. Multiple coordinated experimental efforts, involving large-scale trial-and-error methods, are required to investigate new compounds. In general, this is due to the inherent difficulties in identifying novel therapeutic targets such as genes that cause disease. Even where potential target genes are identified robustly, it is difficult to find drug candidate compounds that successfully bind to the proteins they encode.

Computer-based methods have been introduced in an attempt to shorten the period of drug development and to reduce the expenses involved. The two major computer-aided drug design strategies are ligand-based drug design (LBDD) and structure-based drug design (SBDD). LBDD has various advantages including less required computational resources and better success rates for drug design. However, it also has the disadvantage of limited ability to find drug candidate compounds with low structural similarity to known drugs. To compensate for the weaknesses of LBDD, SBDD shows a greater ability to find drug candidate compounds lacking in structural similarity with known drugs. This is because SBDD tries to screen drug candidate compounds by investigating whether these can bind to target proteins. The weak point of SBDD is that it requires massive computational resources, and this prevents its application to large-scale screening, in which candidate drug compounds often number several million.

Considering the relatively low cost of obtaining gene expression profiles, a third computer-aided strategy has been proposed: gene expression profile-based drug design. In this strategy, the gene expression profiles of tissues/cell lines treated with candidate drug compounds are collected. The collected profiles are then compared with those of tissues/cell lines treated with known drug compounds. If the candidate drug compounds share a gene expression profile to some extent with known drug compounds, they are identified as having therapeutic potential against target diseases/proteins.

Some databases have been established to assist gene expression profiling for drug design. For example, chemical checker (10) includes gene expression in computer-aided drug design, whereas PharmacoDB (49) is fully implemented to consider the dose dependence of drug-treated cell lines for drug design. Many papers have been published on the use of gene expression profiles for computer-aided drug design (8, 3). For instance, Huang et al (18) used combinatorial analysis of drug-induced gene expression for cancer drugs, which were then experimentally confirmed *in vitro*. Lee et al. (31) proposed DeSigN, a robust and useful method for identifying candidate drugs using an input gene signature obtained from gene expression analysis. Kim et al. (25) performed computational drug repositioning for gastric cancer using reversal of gene expression profiles, and Wolf et al. (9) analyzed high-throughput gene expression profiles to identify similarities between drugs and to predict compound activity. Hodos et al. (17) tried to fill in missing gene expression observations in cells treated with drugs by predicting cell-specific drug perturbation profiles using available expression data from related conditions. Pabon et al. (40) predicted protein targets for drug-like compounds using transcriptomics. In contrast, Liu et al. (33) performed comparative analysis of genes that are frequently regulated by drugs based on connectivity to map transcriptome data.

In contrast to these successful applications of gene expression profile analysis to computer-aided drug design, it is unclear how individual gene expression is affected by drug treatment. First, the number of genes expressed in a dose dependent-manner is as large as the number of genes expressed. Thus, it is not easy to invent a useful method to integrate and understand the dose dependent-genes pertaining to individual gene expression profiles. For example, Luka et al. (35) employed principal component analysis (PCA) to integrate the dose dependence of gene expression profiles upon combinatorial drug treatment. They reported a convex (not monotonic) dependence on dose density and identified this as evidence of the cooperative effects of dual drug treatments. Nevertheless, convex dependence on dose was reportedly observed in a single drug treatment if tensor decomposition (TD) was employed to integrate multiple gene expression profiles of cell lines treated with a single drug (13). Thus, it is primarily important to identify an effective method that can integrate numerous gene expression profiles of tissues/cell lines treated with drugs.

Recently, Kozawa et al. (28) used the gene expression profiles of mouse tissues treated with drugs to predict human clinical outcomes. In this paper, we applied TD-based unsupervised feature extraction (FE) to the gene expression profiles used in their study and attempted to identify the changes in gene expression profiles of mouse tissues treated with individual drugs.

## METHODS AND MATERIALS

Figure 1 shows the flow chart of analysis.

**Figure 1.**
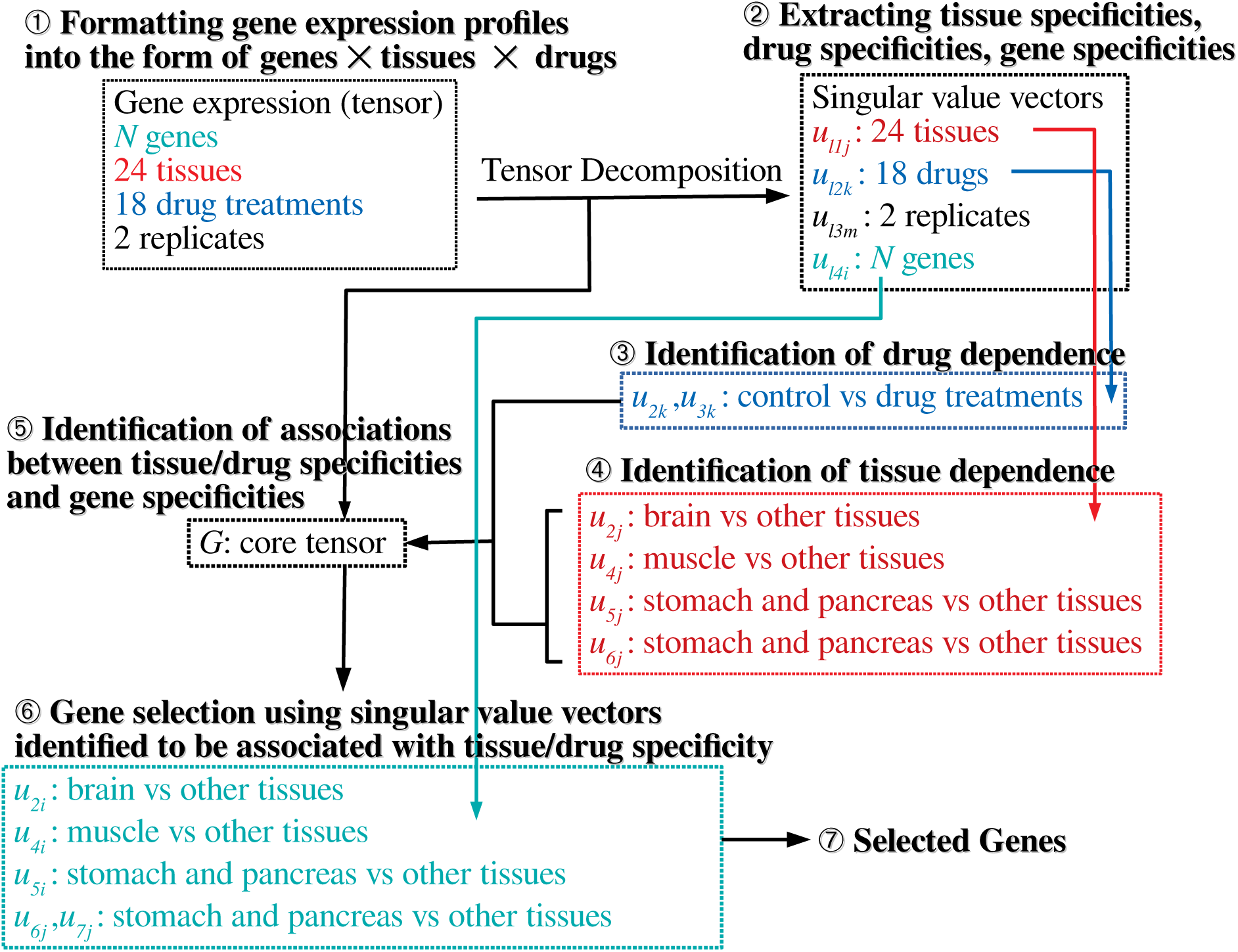
Schematic of the analysis performed in this study

### Gene expression profiles

The gene expression profiles used in this study were downloaded from the gene expression omnibus (GEO) with GEO ID GSE142068. Twenty four profiles named “GSE142068 count XXXXX.txt.gz” were downloaded, where “XXXXX” indicates one of the 24 tissues, i.e., AdrenalG, Aorta, BM (Bone marrow), Brain, Colon, Eye, Heart, Ileum, Jejunum, Kidney, Liver, Lung, Pancreas, ParotidG, PituitaryG, SkMuscle, Skin, Skull, Spleen, Stomach, Spleen, Thymus, ThyroidG, and WAT (white adipose tissue), which were treated with 15 drugs: Alendronate, Acetaminophen, Aripiprazole, Asenapine, Cisplatin, Clozapine, Clozapine, Empagliflozin, Lenalidomide, Lurasidone, Olanzapine, Evolocumab, Risedronate, Sofosbuvir, and Teriparatide, and Wild type (WT).

### TD-based unsupervised FE

For applying TD-based unsupervised FE (14) to the downloaded gene expression profiles, they must be formatted as a tensor. In this analysis, they were formatted as tensor, *x*_*ijkm*_ ∈ ℝ^*N*×24×18×2^, for *N* genes, 24 tissues, 18 drug treatments, and two replicates. Then, the HOSVD (14) algorithm was applied to *x*_*ijkm*_ and we obtained TD

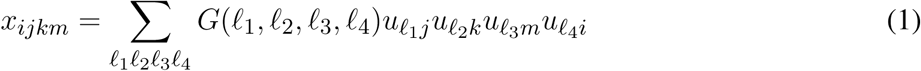

where *G* ∈ ℝ^*N*×24×18×2^ is the core tensor, 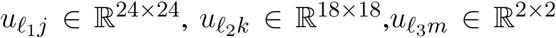, and 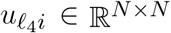, represents singular value matrices that are also orthogonal matrices. *x*_*ijkm*_ is considered to be standardized as ∑_*i*_ *x*_*ijkm*_ = 0 and 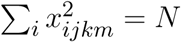.

Mathematically, eq. (1) aims to decompose the dependence of *x*_*ijkm*_ upon *i, j, k, m* into a series of products among 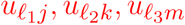, and 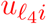, each of which is supposed to represent the dependence on *i, j, k, m*. As it is unlikely that a single product of 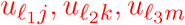, and 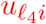 can reproduce *x*_*ijkm*_, we need to consider various combinations of 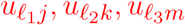 and 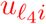, where those associated with distinct *ℓ*_1_, *ℓ*_2_, *ℓ*_3_, *ℓ*_4_ are supposed to be associated with distinct dependence on *i, j, k, m*. Then, the products of 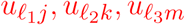, and 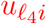, must be summed up with the weight of *G* to reproduce *x*_*ijkm*_. Biologically, we cannot expect that 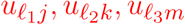, and 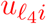 can represent the biological aspect because eq. (1) is simply a mathematical hypothesis; therefore, their association with a biological aspect after performing TD needs to be validated.

To understand how gene expression profiles are altered by drug treatment in a tissue-group-wide manner, we first need to investigate 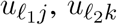, and 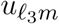 After identifying which *ℓ*_1_, *ℓ*_2_ and *ℓ*_3_ are biologically interesting, we select *ℓ*_4_ associated with *G*(*ℓ*_1_, *ℓ*_2_, *ℓ*_3_, *ℓ*_4_) that have the largest absolute values with fixed *ℓ*_1_, *ℓ*_2_ and *ℓ*_3_, because 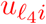 associated with such *ℓ*_4_ is supposed to represent the weight of gene *i* that is expressed in association with *j, k, m* dependence represented by the selected 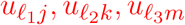.

Using the identified 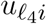, the *P* -value, *P*_*i*_, is attributed to gene *i* as

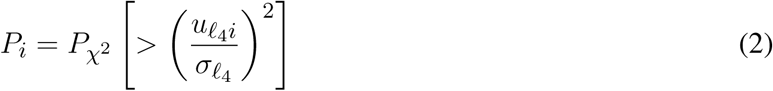

where 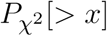 is the cumulative probability of *χ*^2^ distribution and 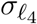 is the standard deviation. Here, we assume that 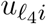 obeys a Gaussian distribution with zero mean because ∑_*i*_ *x*_*ijkm*_ = 0. *P*_*i*_ is corrected via the BH criterion (6) and *I*, a set of genes *i* associated with adjusted *P* -values less than 0.01, is selected. For a more detailed explanation of TD-based unsupervised FE, see the recently published monograph (14). *t* **test and Wilcoxon test applied to sets of genes classified based on tissue groups and drugs groups**

In order to determine whether the selected set of genes, *I*, are expressed distinctly between the two assigned tissue groups, *J*, {*x*_*ijkm*_ |*i* ∈ *I, j* ∈ *J*}, and 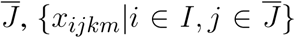, we applied a two-way *t* test and Wilcoxon test and computed the *P* -values. Similar analyses were done with two drug groups, *K*, {*x*_*ijkm*_ |*i* ∈ *I, k* ∈ *K*}, and 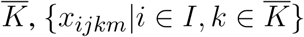.

### Enrichment analysis

The selected genes (gene symbols) were uploaded to Enrichr (30) and Metascape (56) in order to validate the various biological functions of the selected genes.

## RESULTS

Figure 2 summarizes the results obtained in this study.

**Figure 2.**
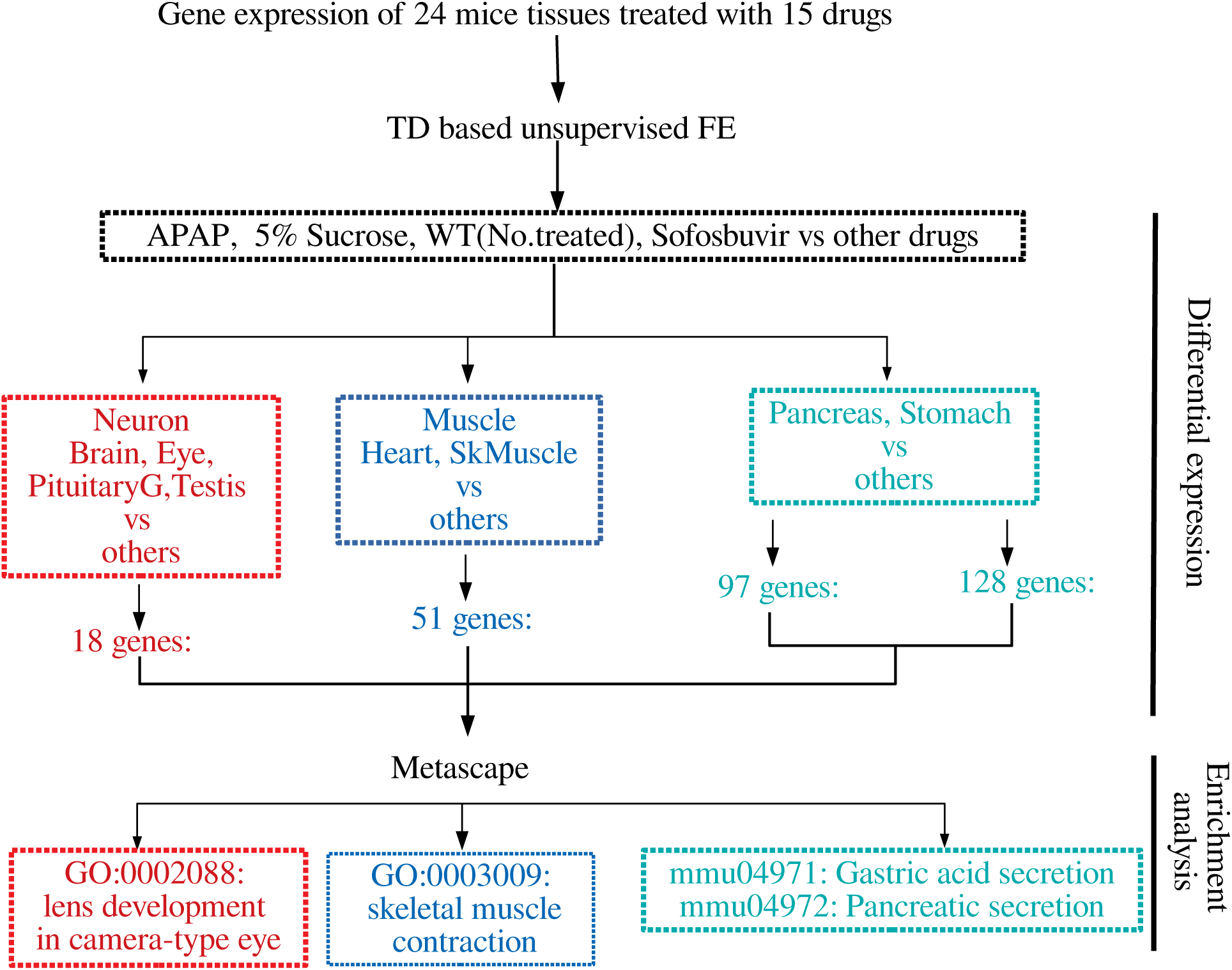
Summary of the results obtained in this study

### Drug treatment specificity

After obtaining the TD, eq. (1), we first investigated 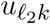 attributed to the *k*th drug. Although the number of drugs tested is as many as 15, the total number of drug treatments was considered to be 18 due to the testing of three additional conditions. Usually, the first singular value vectors represent uniform values (i.e., components that are not distinct between samples) (14). In this case, *u*_1*k*_ does not represent any dependence on drug treatment. This is reasonable because the expression of most genes is unlikely to be affected by drug treatment. We thus considered the second and third singular value vectors, *u*_2*k*_ and *u*_3*k*_, attributed them to drug treatments (Fig. 3). In contrast to expectations, the drug treatments were quite universal. Most of the drug treatments (other than (2), (9), (15), and (17)) were separated from the control treatments ((2), (9), (15), and (17)) along one direction (red arrow) whereas the diversity among drug treatments was spread perpendicular (blue arrow) to that direction, only among drug treatments. This suggests that the gene expression profiles are altered similarly, independently of the drug treatment.

**Figure 3.**
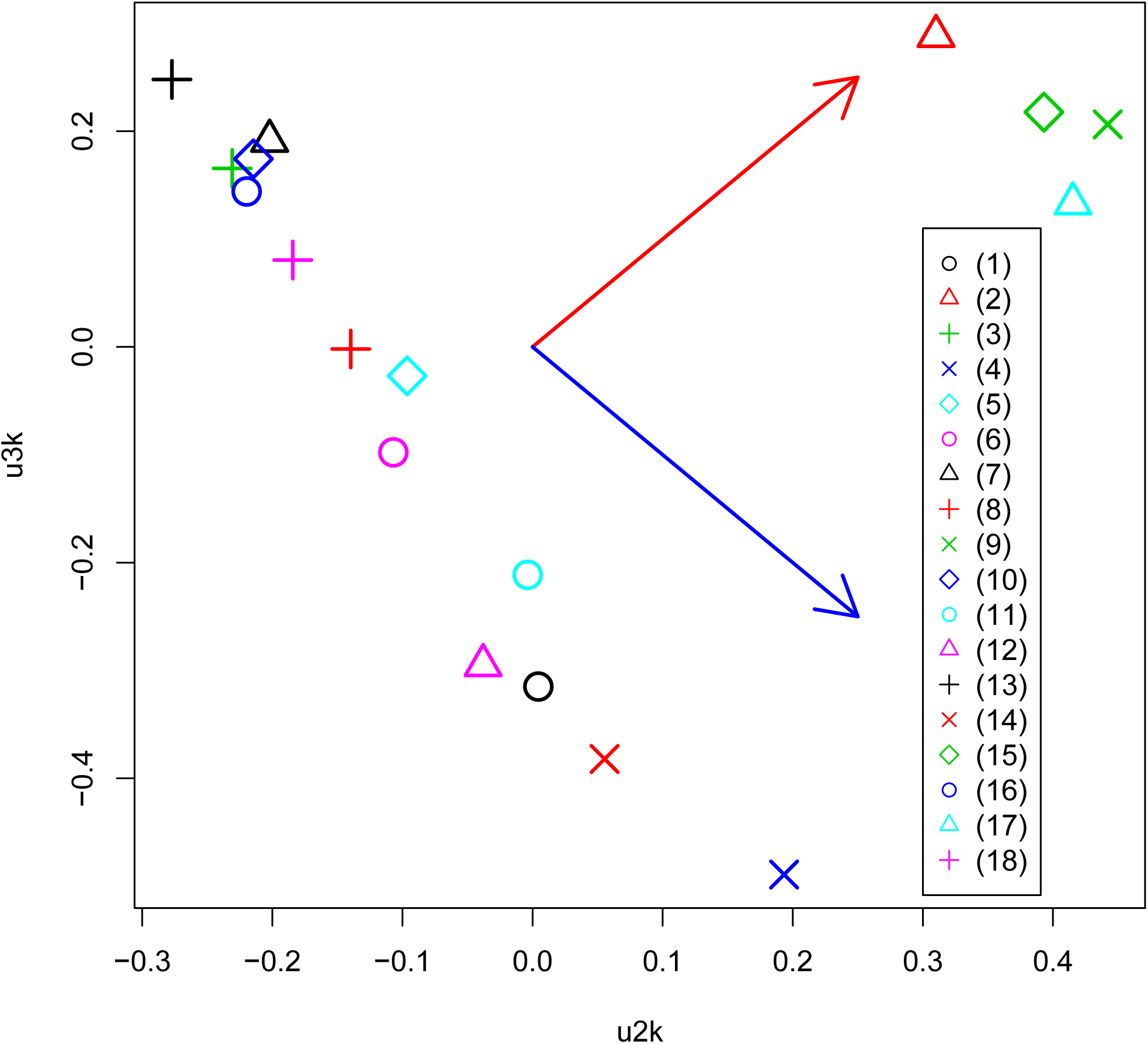
Scatter plot between *u*_2*k*_ and *u*_3*k*_ attributed to drug treatment. Red and blue arrows represent distinct controls and drug treatments, and diversity among drug treatments, respectively. (1) Alendronate, (2) APAP, (3) Aripiprazole, (4) Asenapine, (5) Cisplatin, (6) Clozapine, (7) Dox, (8) EMPA, (9) FivePercentSucrose, (10) Lenalidomide, (11) Lurasidone, (12) Olanzapine, (13) Repatha, (14) Risedronate, (15) Sofosbuvir, (16) Teriparatide, (17) WT.No.treated, (18) 5percentCMC0.25percentTween80.

### Tissue-specificity

We further studied the relationship of universal drug treatments with individual tissues. For this, we next investigated 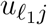 attributed to 24 tissues. We then found that several 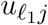 are expressed in a tissue-group wide manner (Fig. 4). The tissue-wide expression pattern identified by singular value vectors is described as follows; As *u*_1*j*_ does not express any tissue specificities, it is unlikely to exhibit tissue specificity; as *u*_2*j*_ has larger absolute values for the brain, eye, pituitary, and testis, it is likely to represent neuronal tissue specificities; as *u*_3*j*_ has larger absolute values only for the parotid, we did not consider it further; as *u*_4*j*_ exhibits larger absolute values for the heart and SkMuscle, we considered that it exhibits muscle specificities; As *u*_5*j*_ and *u*_6*j*_ exhibit larger absolute values for the pancreas and stomach, we considered that it exhibits gastric tissue specificities. It is thus obvious that the combination of tissue specificity is quite reasonable biologically.

**Figure 4.**
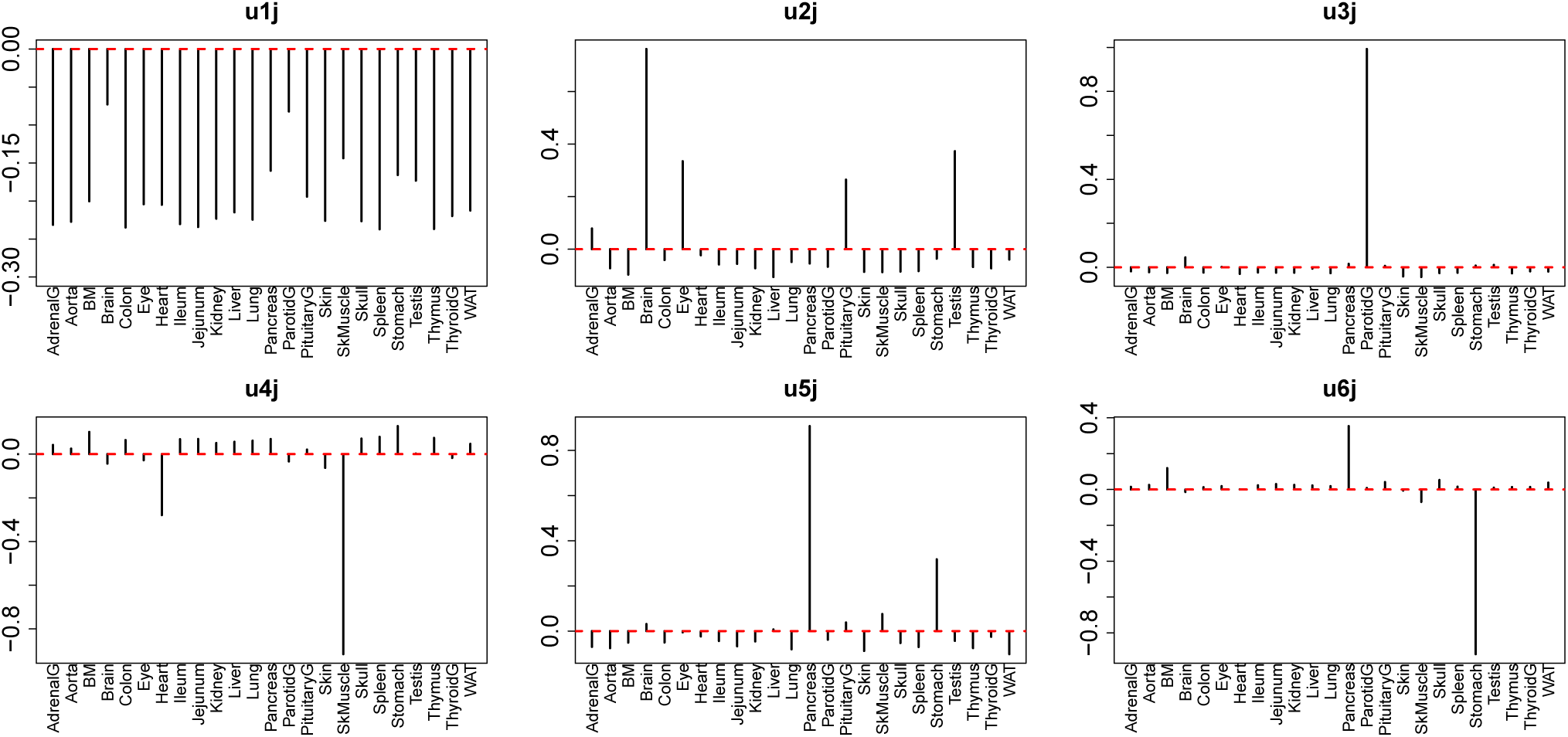
Singular value vectors, 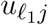, attributed to tissues. *u*_1*j*_: no tissue specificity. *u*_2*j*_: Brain Eye, Pituitary, and Testis, thus mostly neuron-specific. *u*_3*j*_: Parotid-specific, *u*_4*j*_: Heart and SkMuscle, thus muscle-specific, *u*_5*j*_ and *u*_6*j*_: stomach and pancreas, thus, gastrointestinal-specific.

Aiming to specify singular value vectors attributed to genes, 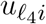, for gene selection, we then checked which of *G*(*ℓ*_1_, 2, 1, *ℓ*_4_) and *G*(*ℓ*_1_, 3, 1, *ℓ*_4_) have larger absolute values, as *u*_1*m*_ always exhibits the same values between two replicates (Table 1).

**Table 1.**
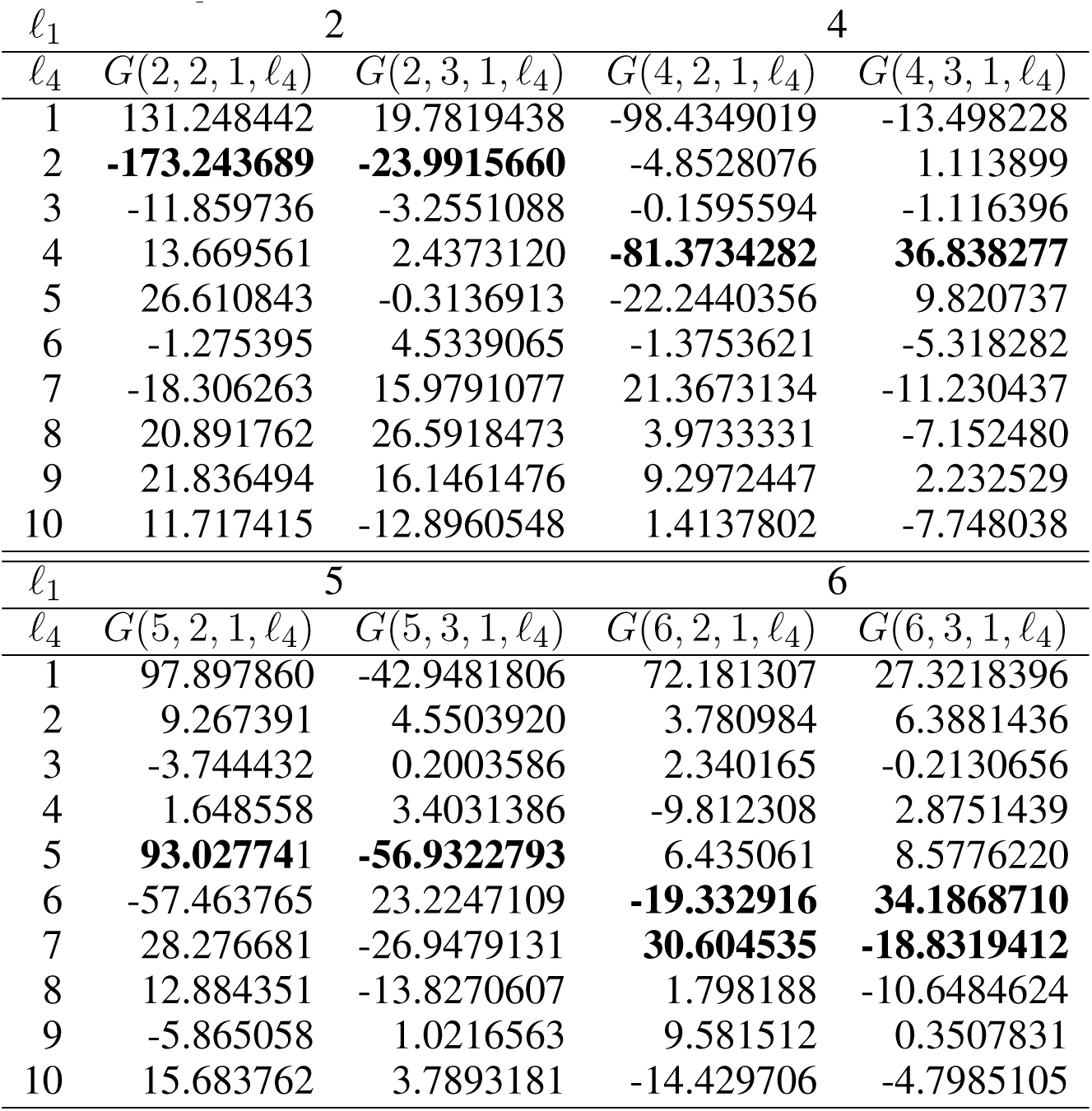
*G*(*ℓ*_1_, 2, 1, *ℓ*_4_) and *G*(*ℓ*_1_, 3, 1, *ℓ*_4_) for *ℓ*_1_ = 2, 4, 5, 6. Values in bold correspond to those of *ℓ*_4_s used for gene selection with 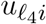.

For *ℓ*_1_ = 2, which is supposed to be attributed to neuron-specific tissues (*u*_2*j*_), *G*s with *ℓ*_4_ = 2 have larger absolute values. Thus, *u*_2*i*_ was employed for neuron-specific gene selection. For *ℓ*_1_ = 4, which is supposed to be attributed to muscle-specific tissues (*u*_4*j*_), *G*s with *ℓ*_4_ = 4 have larger absolute values. Thus, *u*_4*i*_ was employed for muscle-specific gene selection. For *ℓ*_1_ = 5, which is supposed to be attributed to gastrointestinal-specific tissues (*u*_5*j*_), *G*s with *ℓ*_4_ = 5 have larger absolute values. Thus, *u*_5*i*_ was employed for muscle-specific gene selection. For *ℓ*_1_ = 6, which is also supposed to be attributed to gastrointestinal-specific tissues (*u*_6*j*_), *G*s with *ℓ*_4_ = 6, 7 have larger absolute values. Then, *u*_6*i*_ and *u*_7*i*_ were employed for muscle-specific gene selection.

After computing the adjusted *P* -values, *P*_*i*_, attributed to the genes (see methods), the genes associated with adjusted *P*_*i*_ less than 0.01 were selected (Table 2. The lists of selected genes can be found in supporting information (Additional File 1). Figure 5 shows a Venn diagram of the selected genes. As expected, two sets of genes, Gas1 and Gas2, which are supposed to be gastrointestinal-specific, are quite common. Other than these, the selected genes are quite distinct from one another. Thus, TD-based unsupervised FE successfully identified the genes whose expression was affected by the drugs in a tissue group-specific manner.

**Table 2.**
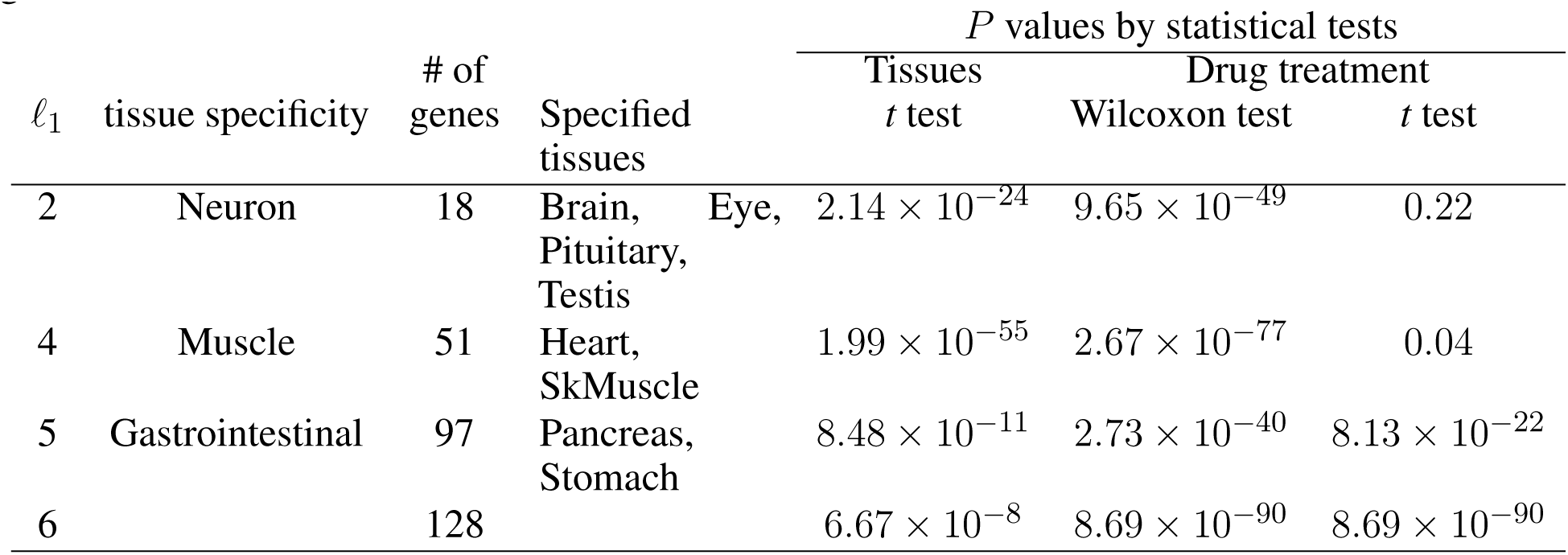
Statistical tests for distinct expression between the specified tissues and other tissues, and between drug treatments and controls.

**Figure 5.**
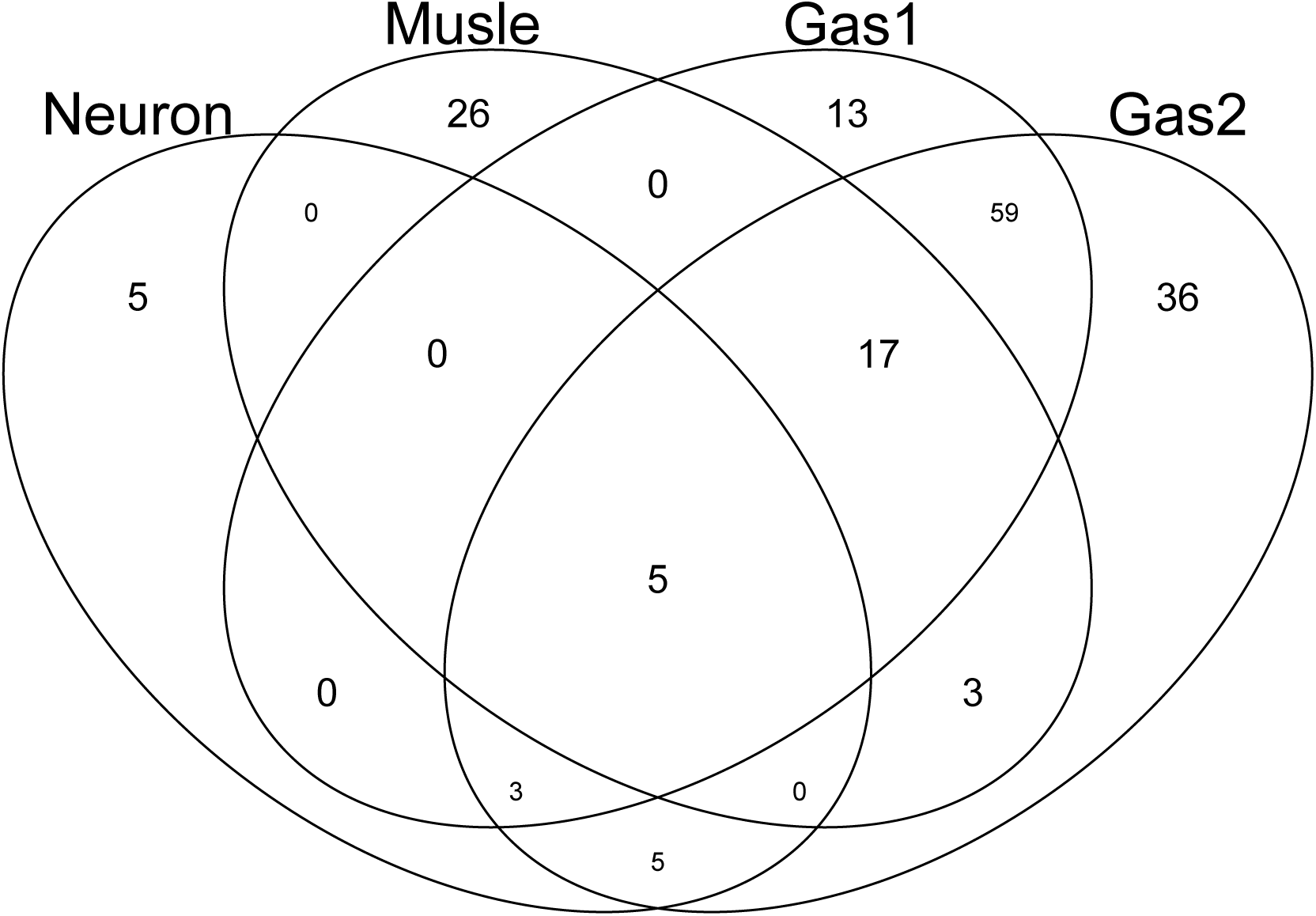
Venn diagram of genes selected by TD-based unsupervised FE. Neuron: genes associated with *u*_2*j*_, which is supposed to be neuron-specific. Muscle: genes associated with *u*_4*j*_, which is supposed to be muscle-specific. Gas1 and Gas2: genes associated with *u*_5*j*_ and *u*_6*j*_ respectively, which is supposed to be gastrointestinal-specific.

### Confirmation of differential expression

In order to check whether the selected genes are expressed distinctly between the specified tissues and other tissues, as well as between drug treatments and controls, we first applied statistical tests to the selected genes (Table 2). The data clearly showed that for all cases, gene expression was distinct between the specified tissues and other tissues as well as between drug treatments and controls. Thus, TD-based unsupervised FE allowed us to select the genes whose expression is coincident with 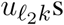 in Fig. 3 and 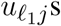 in Fig. 4.

### Biological evaluation

Next, we evaluated the selected genes biologically. For this purpose, we first uploaded the genes to Metascape (Fig. 6). Initially, we noticed that Gas1 and Gas2 largely shared the enriched terms as expected, even though these two gene sets were selected using distinct singular values (*u*_5*i*_ and *u*_6*i*_, *u*_7*i*_, respectively). In particular, it is important to note that two KEGG terms, “mmu04971: Gastric acid secretion” and “mmu04972: Pancreatic secretion” are shared by Gas1 and Gas2, which are supposed to be Pancreas- and Stomach-specific. In contrast, various muscle-related terms are enriched in the Muscle gene set as expected, whereas “GO:0002088: lens development in camera-type eye” is enriched in the neuronal gene set. All of these results suggest that TD-based unsupervised FE selected the biologically reasonable genes.

**Figure 6.**
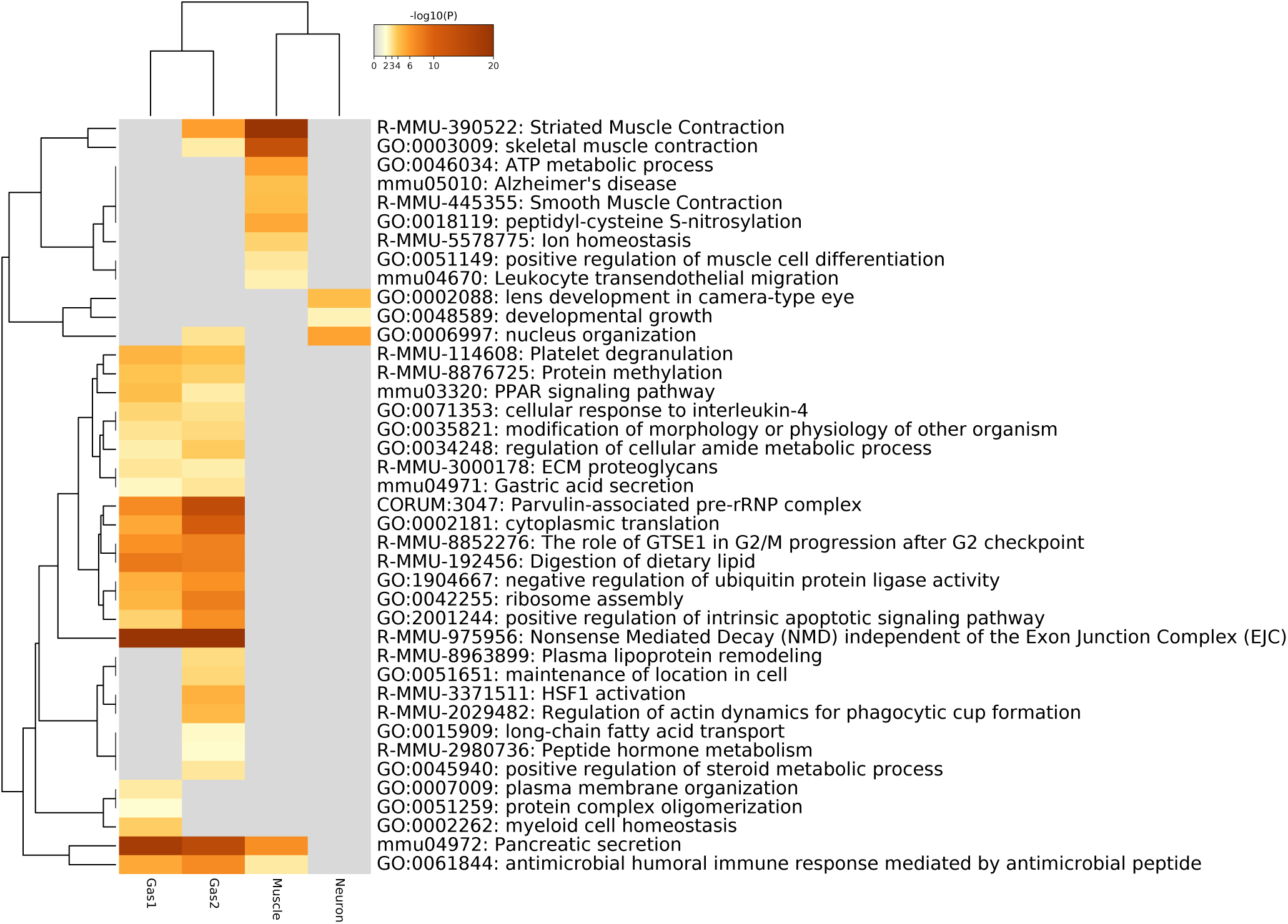
Heatmap of enrichment analysis provided by Metascape.

Figure 7 shows the protein-protein interaction (PPI) network provided by Metascape. A high degree of connectivity was obvious. Thus, TD-based unsupervised FE identified the sets of genes among which PPI is enriched. Moreover, Gas1 and Gas2 largely share the PPI network, whereas the neuronal and muscular gene sets form their own PPI network within which PPI is enriched. Thus, PPI analysis also indicated that TD-based unsupervised FE identified biologically reasonable genes.

**Figure 7.**
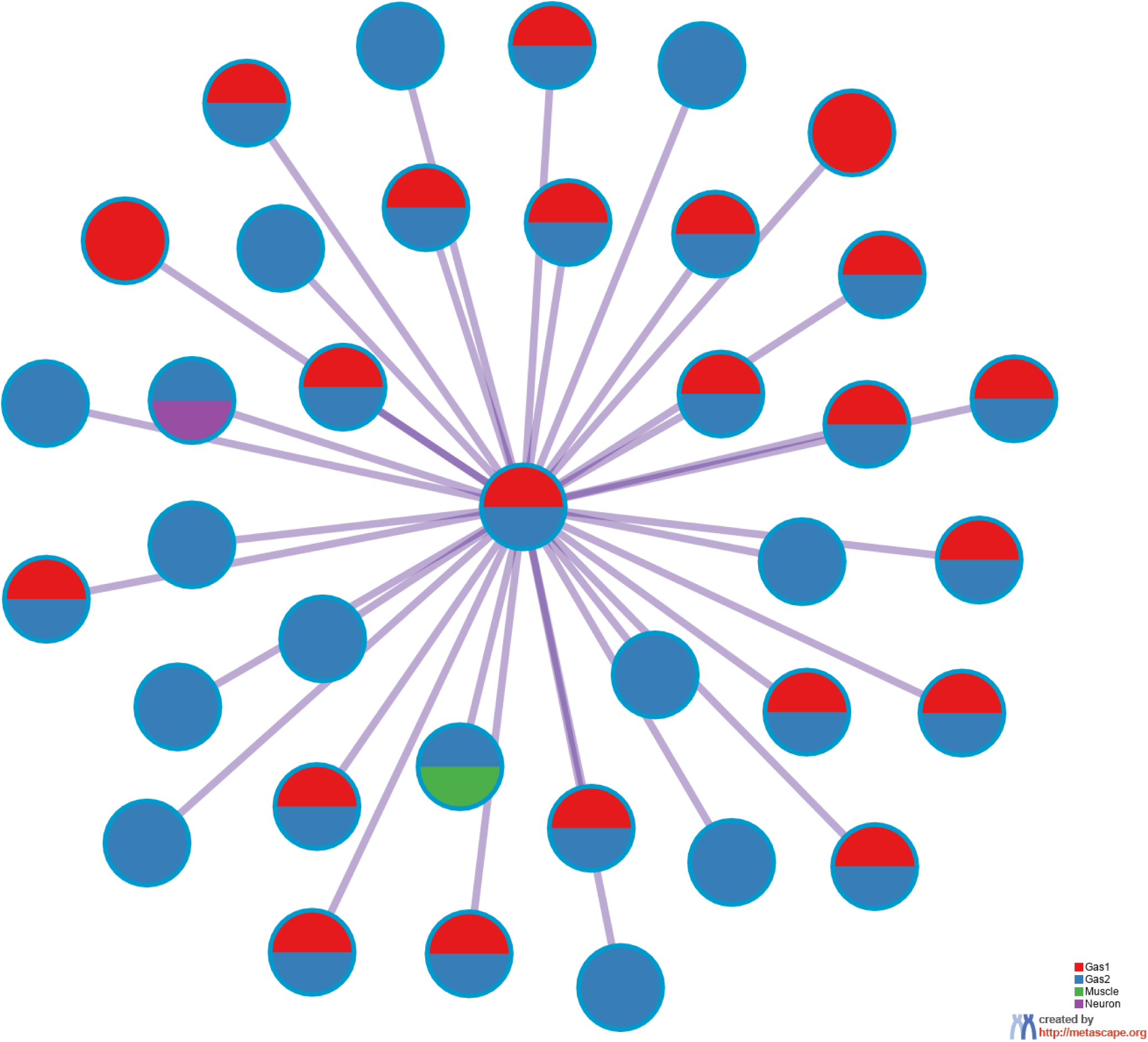
PPI network provided by Metascape. Red: Gas1, Blue: Gas2, Green: Muscle, Purple: Neuron.

To eliminate the possibility that our results were specific to the Metascape data set, we uploaded the genes selected by TD-based unsupervised FE to Enrichr (Table 3). With this data set, we observed clear detection of at least one tissue-related disease for four sets of tissue-specific genes, validating the Metascape-based results.

**Table 3.**
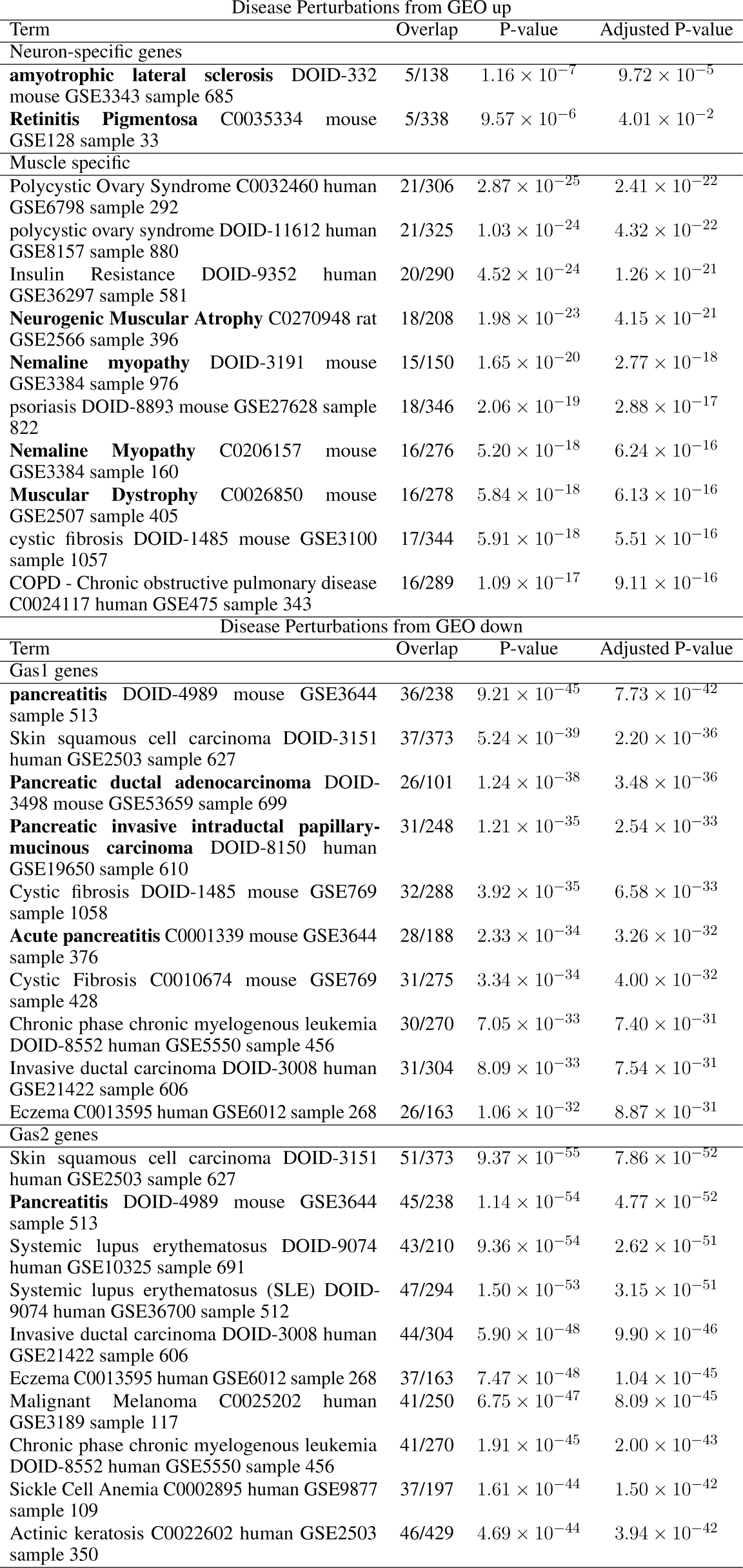
Enrichment analysis for “Disease Perturbations from GEO up” and “Disease Perturbations from GEO down” by Enrichr. Diseases in bold correspond to those related to specific tissues. Up to the top 10 ranked terms are shown.

## DISCUSSION

Although it is unclear why the 15 drugs affected the expression of many common genes, a detailed investigation can allow further interpretation. Table 4 shows the drugs’ effects on neuronal, muscular, and pancreatic tissues. These data suggest that most drugs simultaneously affect these three groups of tissues.

**Table 4.** Previously reported drug effects on neuron (brain and eye), muscle and pancreas tissues. (*): Reported side effects.

| Drugs | Tissue types |  |  |
| --- | --- | --- | --- |
|  | Neuron | Muscle | Pancreas or Stomach |
| Alendronate | Brain calcification (39) | Muscle mass (15) | Pancreatitis (19) |
| Acetaminophen (APAP) | Brain (12) | Skeletal muscle (51) | Pancreatitis (7) |
| Aripiprazole | Brain Activation (37) | Muscle spasms (*) | Pancreatitis (26) |
| Asenapine | Cognitive and monoamine dysfunction (11) | Muscle rigidity(*) | — |
| Cisplatin | Prefrontal cortex (20) | Muscle atrophy (47) | Pancreas (54) |
| Clozapine | Brain (32) | Myotoxicity (45) | Pancreatitis (5) |
| Doxycycline | Brain (34) | Smooth Muscle (4) | Acute pancreatitis (43) |
| Empagliflozin | Neurovascular unit and neuroglia (16) | Muscle sympathetic nerve activity (21) | Pancreatitis (27) |
| Lenalidomide | Memory loss (46) | Muscle cramp (44) | Pancreatic cancer (52) |
| Lurasidone | Acute schizophrenia (55) | Muscle(*) | — |
| Olanzapine | Brain stem (1) | Acute muscle toxicity (24) | Pancreatitis (23) |
| Repatha (Evolocumab) | — | Muscle-related statin Intolerance (38) | — |
| Risedronate (actonel) | Ocular myasthenia (42) | Muscle weakness (2) | Gastrointestinal cancer (53) |
| Sofosbuvir | Ocular surface (48) | Myositis (41) | Pancreatitis (36) |
| Teriparatide | — | Muscle Cramp (22) | Pancreatitis (*) |

Our results are in contrast to the study that inspired our work (28), in which the authors employed a fully supervised approach requiring previous knowledge. Although Kosawa et al. (28) also aimed to infer the therapeutic and side effects of drug treatments in humans based on gene expression in drug-treated tissues, their analysis required previous knowledge that is not needed for TD-based unsupervised FE. In this sense, our approach has distinct potential that the original study could not achieve.

In addition to the above-mentioned biological superiority of TD-based unsupervised FE, this approach also has some methodological advantages as follows. First, although we classified 24 tissues into two groups based on the observation of singular value vectors attributed to tissues, 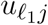 (Fig. 4) prior to the identification of differentially expressed genes, it is computationally infeasible for other methods to classify 24 tissues into two groups before starting to seek differentially expressed genes, as there are no criteria on how to divide 24 tissues into two groups. It is thus practically impossible to analyze all possible divisions, as they number in the millions. The same advantage is observed when grouping 18 drug treatments into two. This may be much easier than classifying tissues, because some of the drug treatments are obviously controls. Nevertheless, based upon the second and third singular value vectors attributed to drug treatments, *u*_2*k*_ and *u*_3*k*_ (Fig. 3), acetaminophen (APAP) and sofosbuvir are grouped together with two control treatments. Such a classification can never be proposed without TD. In this sense, there is no computationally feasible method that can compete with our method.

The biological basis for the two groups of drugs seen in Fig. 3 may be questioned. To clarify this point, we uploaded two groups of drugs to DrugEnrichr (29), which evaluates the coincidence of genes targeted by the uploaded drugs (Additional file 2). Based on the “Geneshot Predicted from Co-expression” category in DrugEnrichr, we found that there are at least as many as 164 genes targeted by two drugs (APAP and Sofosbuvir) in group1 whereas 213 genes are targeted by at least two drugs among as many as thirteen drugs included in group2 (Alendronate, Aripiprazole, Asenapine, Cisplatin, Clozapine, Dox, EMPA, Lenalidomide, Lurasidone, Olanzapine, Repatha, Risedronate, Teriparatide). This suggests that these two groups of drugs are quite distinct because there are no common targeted genes between these 164 and 213 genes. Thus, the groups of drugs identified by TD based unsupervised FE are likely based on the genes that the drugs target.

In view of the two above-mentioned advantages, TD-based unsupervised FE might yield completely distinct outcomes that other supervised methods cannot, and it therefore represents a worthwhile primary or supplementary approach to gene-expression-based investigation of drug effects.

One might wonder if the results were confirmed only by single experiments. As the results shown in Table 3 indicate coincidence between the present result and other studies, the results derived in this study are not dependent on a single study, but are coincident with numerous studies in the public domain database.

Moreover, TD-based unsupervised FE is a very useful strategy for repositioning known drugs. As shown in Fig. 3, TD-based unsupervised FE can determine the effective tissue. Furthermore, as indicated in Table 3, the genes selected by TD-based unsupervised FE can indicate the diseases for which the drugs have potential effectiveness. Therefore, applying TD-based unsupervised FE to gene expression profiles altered by drug treatments can be a promising strategy to repurpose known drugs for new diseases.

## CONCLUSIONS

In this paper, we applied TD-based unsupervised FE (14) to the gene expression profiles of 24 mouse tissues treated with 15 drugs. Integrated analysis allowed us to identify the universal nature of drug treatments in a tissue-group-wide manner, which is generally impossible to identify using any other supervised strategy that requires prior information.

## Supporting information

Genes selected by TD based unsuoervised FE

Genes predicted by DrugEnrichr

## DATA AVAILABILITY

All datasets analyzed in this study were obtained from the GEO database.

## AUTHOR CONTRIBUTIONS

YHT planned and performed the study. YHT and TT discussed the results and wrote the paper.

## ACKNOWLEDGMENTS

The authors would like to thank the reviewers for very constructive comments and thoughtful suggestions. This manuscript has been released as a pre-print at BioRxiv (Taguchi Y-h. et al, 2020 (50)).

## ADDITIONAL FILES

**Additional file 1 — List of genes selected by TD-based unsupervised FE**

List of genes shown in Table 2 (xlsx)

**Additional file 2 — List of genes predicted by DrugEnrichr, whose expression is likely affected by the drugs investigated in this study**

List of genes (xlsx)

## REFERENCES

1. Anwar, I. J., Miyata, K., and Zsombok, A. (2016). Brain stem as a target site for the metabolic side effects of olanzapine. Journal of Neurophysiology 115, 1389–1398. doi: 10.1152/jn.00387.2015. PMID: 26719086

2. Badayan, I. and Cudkowicz, M. E. (2009). Profound muscle weakness and pain after one dose of actonel. Case Reports in Medicine 2009, 1–3. doi: 10.1155/2009/693014

3. Bates, S. (2011). The role of gene expression profiling in drug discovery. Current Opinion in Pharmacology 11, 549 – 556. doi: https://doi.org/10.1016/j.coph.2011.06.009. Anti-infectives/New technologies

4. Bendeck, M. P., Conte, M., Zhang, M., Nili, N., Strauss, B. H., and Farwell, S. M. (2002). Doxycycline modulates smooth muscle cell growth, migration, and matrix remodeling after arterial injury. The American Journal of Pathology 160, 1089–1095. doi: 10.1016/s0002-9440(10)64929-2

5. Bergemann, N., Ehrig, C., Diebold, K., Mundt, C., and v. Einsiedel, R. (1999). Asymptomatic pancreatitis associated with clozapine. Pharmacopsychiatry 32, 78–80. doi: 10.1055/s-2007-979197

6. Burgos, K., Malenica, I., Metpally, R., Courtright, A., Rakela, B., Beach, T., et al. (2014). Profiles of extracellular mirna in cerebrospinal fluid and serum from patients with alzheimer’s and parkinson’s diseases correlate with disease status and features of pathology. PLOS ONE 9, 1–20. doi: 10.1371/journal.pone.0094839

7. Chen, S.-J., Lin, C.-S., Hsu, C.-W., Lin, C.-L., and Kao, C.-H. (2015). Acetaminophen poisoning and risk of acute pancreatitis. Medicine 94, e1195. doi: 10.1097/md.0000000000001195

8. Chengalvala, M. V.,, Chennathukuzhi, V. M., Johnston, D. S., Stevis, P. E., and Kopf, G. S. (2007). Gene expression profiling and its practice in drug development. Current Genomics 8, 262–270. doi: 10.2174/138920207781386942

9. De Wolf, H., Cougnaud, L., Van Hoorde, K., De Bondt, A., Wegner, J. K., Ceulemans, H., et al. (2018). High-throughput gene expression profiles to define drug similarity and predict compound activity. ASSAY and Drug Development Technologies 16, 162–176. doi: 10.1089/adt.2018.845. PMID: 29658791

10. Duran-Frigola, M., Pauls, E., Guitart-Pla, O., Bertoni, M., Alcalde, V., Amat, D., et al. (2020). Extending the small molecule similarity principle to all levels of biology. Nature Biotechnology doi: 10.1101/745703. In press

11. Elsworth, J. D., Groman, S. M., Jentsch, J. D., Valles, R., Shahid, M., Wong, E., et al. (2012). Asenapine effects on cognitive and monoamine dysfunction elicited by subchronic phencyclidine administration. Neuropharmacology 62, 1442 – 1452. doi: https://doi.org/10.1016/j.neuropharm.2011.08.026.Schizophrenia

12. Ghanem, C. I., Pérez, M. J., Manautou, J., and Mottino, A. D. (2016). Acetaminophen from liver to brain: New insights into drug pharmacological action and toxicity. Pharmacological Research 109, 119 – 131. doi: https://doi.org/10.1016/j.phrs.2016.02.020. Country in focus: Pharmacology in Argentina

13. h. Taguchi, Y. (2019). Drug candidate identification based on gene expression of treated cells using tensor decomposition-based unsupervised feature extraction for large-scale data. BMC Bioinformatics 19. doi: 10.1186/s12859-018-2395-8

14. Taguchi, Y. (2020). Unsupervised Feature Extraction Applied to Bioinformatics: A PCA Based and TD Based Approach (Switzerland: Springer International Publishing). doi: 10.1007/978-3-030-22456-1

15. Harada, A., Ito, S., Matsui, Y., Sakai, Y., Takemura, M., Tokuda, H., et al. (2015). Effect of alendronate on muscle mass: Investigation in patients with osteoporosis. Osteoporosis and Sarcopenia 1, 53 – 58. doi: https://doi.org/10.1016/j.afos.2015.07.005

16. Hayden, M. R., Grant, D. G., Aroor, A. R., and DeMarco, V. G. (2019). Empagliflozin ameliorates type 2 diabetes-induced ultrastructural remodeling of the neurovascular unit and neuroglia in the female db/db mouse. Brain Sciences 9. doi: 10.3390/brainsci9030057

17. Hodos, R., Zhang, P., Lee, H.-C., Duan, Q., Wang, Z., Clark, N. R., et al. (2017). Cell-specific prediction and application of drug-induced gene expression profiles. In Biocomputing 2018 (WORLD\ SCIENTIFIC). doi: 10.1142/9789813235533\_0004

18. Huang, C.-T., Hsieh, C.-H., Chung, Y.-H., Oyang, Y.-J., Huang, H.-C., and Juan, H.-F. (2019). Perturbational gene-expression signatures for combinatorial drug discovery. iScience 15, 291 – 306. doi: https://doi.org/10.1016/j.isci.2019.04.039

19. Hung, W. Y. (2014). Contemporary review of drug-induced pancreatitis: A different perspective. World Journal of Gastrointestinal Pathophysiology 5, 405. doi: 10.4291/wjgp.v5.i4.405

20. Huo, X., Reyes, T. M., Heijnen, C. J., and Kavelaars, A. (2018). Cisplatin treatment induces attention deficits and impairs synaptic integrity in the prefrontal cortex in mice. Scientific Reports 8. doi: 10.1038/s41598-018-35919-x

21. Jordan, J., Tank, J., Heusser, K., Heise, T., Wanner, C., Heer, M., et al. (2017). The effect of empagliflozin on muscle sympathetic nerve activity in patients with type ii diabetes mellitus. Journal of the American Society of Hypertension 11, 604 – 612. doi: https://doi.org/10.1016/j.jash.2017.07.005

22. Kakaria, P. J., Nashel, D. J., and Nylen, E. S. (2005). Debilitating Muscle Cramps after Teriparatide Therapy. Annals of Internal Medicine 142, 310–310. doi: 10.7326/0003-4819-142-4-200502150-00023

23. Kerr, T. A., Jonnalagadda, S., Prakash, C., and Azar, R. (2007). Pancreatitis following olanzapine therapy: A report of three cases. Case Reports in Gastroenterology 1, 15–20. doi: 10.1159/000104222

24. Keyal, N., Shrestha, G., Pradhan, S., Maharjan, R., Acharya, S., and Marhatta, M. (2017). Olanzapine overdose presenting with acute muscle toxicity. International Journal of Critical Illness and Injury Science 7, 69–71. doi: 10.4103/2229-5151.201962

25. Kim, I.-W., Jang, H., Kim, J. H., Kim, M. G., Kim, S., and Oh, J. M. (2019). Computational drug repositioning for gastric cancer using reversal gene expression profiles. Scientific Reports 9. doi: 10.1038/s41598-019-39228-9

26. Kiraly, B. and Gunning, K. (2008). A case of pancreatitis associated with aripiprazole in the absence of hyperglycemia. The Primary Care Companion to The Journal of Clinical Psychiatry 10, 484–485. doi: 10.4088/pcc.v10n0612e

27. Kishimoto, M., Yamaoki, K., and Adachi, M. (2019). Combination therapy with empagliflozin and insulin results in successful glycemic control: A case report of uncontrolled diabetes caused by autoimmune pancreatitis and subsequent steroid treatment. Case Reports in Endocrinology 2019, 1–8. doi: 10.1155/2019/9415347

28. Kozawa, S., Sagawa, F., Endo, S., Almeida, G. M. D., Mitsuishi, Y., and Sato, T. N. (2020). Predicting human clinical outcomes using mouse multi-organ transcriptome. iScience, 100791 doi: 10.1016/j.isci.2019.100791

29. [Dataset] Kuleshov, M., Kropiwnicki, E., and Ma’ayan, A. (2020). Drugenrichr

30. Kuleshov, M. V., Jones, M. R., Rouillard, A. D., Fernandez, N. F., Duan, Q., Wang, Z., et al. (2016). Enrichr: a comprehensive gene set enrichment analysis web server 2016 update. Nucleic Acids Research 44, W90–W97. doi: 10.1093/nar/gkw377

31. Lee, B. K. B., Tiong, K. H., Chang, J. K., Liew, C. S., Rahman, Z. A. A., Tan, A. C., et al. (2017). DeSigN: connecting gene expression with therapeutics for drug repurposing and development. BMC Genomics 18. doi: 10.1186/s12864-016-3260-7

32. Li, C. H., Stratford, R. E., de Mendizabal, N. V., Cremers, T. I., Pollock, B. G., Mulsant, B. H., et al. (2014). Prediction of brain clozapine and norclozapine concentrations in humans from a scaled pharmacokinetic model for rat brain and plasma pharmacokinetics. Journal of Translational Medicine 12. doi: 10.1186/1479-5876-12-203

33. Liu, X., Zeng, P., Cui, Q., and Zhou, Y. (2017). Comparative analysis of genes frequently regulated by drugs based on connectivity map transcriptome data. PLOS ONE 12, 1–13. doi: 10.1371/journal.pone.0179037

34. Lucchetti, J., Fracasso, C., Balducci, C., Passoni, A., Forloni, G., Salmona, M., et al. (2018). Plasma and brain concentrations of doxycycline after single and repeated doses in wild-type and app23 mice. Journal of Pharmacology and Experimental Therapeutics doi: 10.1124/jpet.118.252064

35. Lukačišin, M. and Bollenbach, T. (2019). Emergent gene expression responses to drug combinations predict higher-order drug interactions. Cell Systems 9, 423–433.e3. doi: 10.1016/j.cels.2019.10.004

36. Margapuri, J. and Jubbal, S. (2019). 902: Acute pancreatitis following treatment with ledipasvir/sofosbuvir for hepatitis virus infection. Critical Care Medicine 47, 430. doi: 10.1097/01.ccm.0000551651.45733.80

37. Myrick, H., Li, X., Randall, P. K., Henderson, S., Voronin, K., and Anton, R. F. (2010). The effect of aripiprazole on cue-induced brain activation and drinking parameters in alcoholics. Journal of Clinical Psychopharmacology 30, 365–372. doi: 10.1097/jcp.0b013e3181e75cff

38. Nissen, S. E., Stroes, E., Dent-Acosta, R. E., and for the GAUSS-3 Investigators (2016). Efficacy and Tolerability of Evolocumab vs Ezetimibe in Patients With Muscle-Related Statin Intolerance: The GAUSS-3 Randomized Clinical Trial. JAMA 315, 1580–1590. doi: 10.1001/jama.2016.3608

39. Oliveira, J. R. M. and Oliveira, M. F. (2016). Primary brain calcification in patients undergoing treatment with the biphosphanate alendronate. Scientific Reports 6. doi: 10.1038/srep22961

40. Pabon, N. A., Xia, Y., Estabrooks, S. K., Ye, Z., Herbrand, A. K., Evelyn, S., et al. (2018). Predicting protein targets for drug-like compounds using transcriptomics. PLOS Computational Biology 14, 1–24. doi: 10.1371/journal.pcbi.1006651

41. Patel, S., Trakroo, S., Sanaka, S., and Qureshi, K. (2015). Severe myositis with the use of sofosbuvir/ledipasvir for hepatitis c infection: A case of unexpected interactions. American Journal of Gastroenterology 110, S333. doi: 10.14309/00000434-201510001-00761

42. Raja, V., Sandanshiv, P., and Neugebauer, M. (2007). Risedronate induced transient ocular myasthenia. Journal of Postgraduate Medicine 53, 274. doi: 10.4103/0022-3859.37525

43. Rawla, P. and Raj, J. P. (2017). Doxycycline-induced acute pancreatitis: A rare adverse event. Gastroenterology Research 10

44. Reece, D., Kouroukis, C. T., LeBlanc, R., Sebag, M., Song, K., and Ashkenas, J. (2012). Practical approaches to the use of lenalidomide in multiple myeloma: A canadian consensus. Advances in Hematology 2012, 1–14. doi: 10.1155/2012/621958

45. Reznik, I., Volchek, L., Mester, R., Kotler, M., Sarova-Pinhas, I., Spivak, B., et al. (2000). Myotoxicity and neurotoxicity during clozapine treatment. Clinical Neuropharmacology 23, 276–280. doi: 10.1097/00002826-200009000-00007

46. Rollin-Sillaire, A., Delbeuck, X., Pollet, M., Mackowiak, M.-A., Lenfant, P., Noel, M.-P., et al. (2013). Memory loss during lenalidomide treatment: a report on two cases. BMC Pharmacology and Toxicology 14. doi: 10.1186/2050-6511-14-41

47. Sakai, H., Sagara, A., Arakawa, K., Sugiyama, R., Hirosaki, A., Takase, K., et al. (2014). Mechanisms of cisplatin-induced muscle atrophy. Toxicology and Applied Pharmacology 278, 190 – 199. doi: https://doi.org/10.1016/j.taap.2014.05.001

48. Salman, A. G. (2016). Ocular surface changes with sofosbuvir in egyptian patients with hepatitis c virus infection. Cornea 35, 323–328. doi: 10.1097/ico.0000000000000736

49. Smirnov, P., Kofia, V., Maru, A., Freeman, M., Ho, C., El-Hachem, N., et al. (2017). PharmacoDB: an integrative database for mining in vitro anticancer drug screening studies. Nucleic Acids Research 46, D994–D1002. doi: 10.1093/nar/gkx911

50. Taguchi, Y.-h. and Turki, T. (2020). Universal nature of drug treatment responses of drug-tissue-wide model-animal experiments. bioRxiv doi: 10.1101/2020.03.08.982405

51. Trappe, T. A., Carroll, C. C., Dickinson, J. M., LeMoine, J. K., Haus, J. M., Sullivan, B. E., et al. (2011). Influence of acetaminophen and ibuprofen on skeletal muscle adaptations to resistance exercise in older adults. American Journal of Physiology-Regulatory, Integrative and Comparative Physiology 300, R655–R662. doi: 10.1152/ajpregu.00611.2010. PMID: 21160058

52. Ullenhag, G. J., Mozaffari, F., Broberg, M., Mellstedt, H., and Liljefors, M. (2017). Clinical and immune effects of lenalidomide in combination with gemcitabine in patients with advanced pancreatic cancer. PLOS ONE 12, 1–19. doi: 10.1371/journal.pone.0169736

53. Vinogradova, Y., Coupland, C., and Hippisley-Cox, J. (2013). Exposure to bisphosphonates and risk of common non-gastrointestinal cancers: series of nested case–control studies using two primary-care databases. British Journal of Cancer 109, 795–806. doi: 10.1038/bjc.2013.383

54. Yadav, Y. C. (2019). Effect of cisplatin on pancreas and testies in wistar rats: biochemical parameters and histology. Heliyon 5, e02247. doi: 10.1016/j.heliyon.2019.e02247

55. Yasui-Furukori, N. (2012). Update on the development of lurasidone as a treatment for patients with acute schizophrenia. Drug Design, Development and Therapy, 107 doi: 10.2147/dddt.s11180

56. Zhou, Y., Zhou, B., Pache, L., Chang, M., Khodabakhshi, A. H., Tanaseichuk, O., et al. (2019). Metascape provides a biologist-oriented resource for the analysis of systems-level datasets. Nature Communications 10. doi: 10.1038/s41467-019-09234-6

